# Vaccine strains of Rift Valley fever virus exhibit attenuation at the maternal-fetal placental interface

**DOI:** 10.1101/2024.05.31.596800

**Authors:** Cynthia M. McMillen, Christina Megli, Rebecca Radisic, Lauren B. Skvarca, Ryan M. Hoehl, Devin A. Boyles, Jackson J. McGaughey, Brian H. Bird, Anita K. McElroy, Amy L. Hartman

## Abstract

Rift Valley fever virus (RVFV) infection causes abortions in ruminant livestock and is associated with an increased likelihood of miscarriages in women. Using sheep and human placenta explant cultures, we sought to identify tissues at the maternal-fetal interface targeted by RVFV. Sheep villi and fetal membranes were highly permissive to RVFV infection resulting in markedly higher virus titers than human cultures. Sheep cultures were most permissive to wild-type RVFV and ΔNSm infection, while live attenuated RVFV vaccines (LAVs; MP-12, ΔNSs, and ΔNSs/ΔNSm) exhibited reduced replication. The human fetal membrane restricted wild-type and LAV replication, and when infection occurred, it was prominent in the maternal-facing side. Type-I and type-III interferons were induced in human villi exposed to LAVs lacking the NSs protein. This study supports the use of sheep and human placenta explants to understand vertical transmission of RVFV in mammals and whether LAVs are attenuated at the maternal-fetal interface.

**Teaser:** Vaccine strains of Rift Valley fever virus have reduced infection and replication capacity in mammalian placenta

## Introduction

Rift Valley fever (RVF) is a zoonotic, mosquito-transmitted disease in Africa and the Middle East that primarily affects domesticated livestock (sheep, goats, cattle, and camels) and can cause mild to severe disease in humans. Rainy seasons, when mosquito populations expand, are accompanied by high rates of vertical transmission in livestock, leading to fetal death. An entire generation of sheep can be lost in a single outbreak in part due to these “abortion storms”, causing devastating economic impacts on affected regions (1). Abortions in livestock are associated with diffuse placental hemorrhage and necrosis (2) along with cerebral and musculoskeletal deformities in fetuses (3, 4). Given its known effects in pregnant animals, Rift Valley fever virus (RVFV) infection might also cause miscarriages in women (5). A study in Sudan showed that women infected with RVFV during pregnancy are approximately four times more likely to have a second or third trimester miscarriage. Cases of late gestation fetal infection, including infants born at full-term with RVF disease, have been documented in humans (6, 7).

Live attenuated RVFV vaccines (LAVs) are currently used in endemic areas to minimize the incidence of disease in livestock (8, 9); however, the residual virulence of some of these attenuated strains may still lead to adverse events in pregnant animals, including fetal loss or deformity (10, 11). Attenuated vaccine strains of RVFV were generated through serial passaging in vivo, chemical mutagenesis, or by mutating or removing one or both virulence factors, NSs or NSm. Two LAV strains, Smithburn and Clone 13 (similar to ΔNSs), are regularly used in livestock throughout Africa (8, 9). Both strains retain some virulence which can lead to fetal death or teratogenic events in pregnant livestock; therefore, pregnant animals are not routinely vaccinated, leaving them vulnerable to infection by virulent field strains of RVFV (10, 11). Other LAVs were developed to overcome the limitations of Smithburn and Clone-13, including MP-12, an attenuated strain generated by chemical mutagenesis that resulted in nine amino acid mutations across all three segments of the RNA genome (12, 13). MP-12 has undergone early stage clinical trials in humans (14, 15); however, there are conflicting reports on the safety of MP-12 during pregnancy. Sheep vaccinated with high doses of MP-12 during early gestation delivered deformed fetuses (16), whereas those vaccinated with a lower dose during late gestation delivered healthy lambs (17, 18). A next-generation LAV strain made using reverse genetics to delete both NSs and NSm (ΔNSs/ΔNSm) appeared to be safe in pregnant sheep, even when vaccinated during early gestation (19). Although there are no approved RVFV vaccines for humans, clinical trials are underway with updated versions of the ΔNSs/ΔNSm and several other promising candidates (20-22). Identifying a safe and effective vaccine to protect humans from severe RVF will be a tremendous benefit to the field. However, early stage human clinical trials are unlikely to provide sufficient safety data in pregnant women as they are generally excluded from these trials. Moreover, as RVFV is particularly pathogenic to pregnant mammals, it is critical to understand the pathogenic mechanisms at the maternal-fetal interface.

Despite the well-documented outcomes of RVFV fetal infection in ruminant livestock, it is not clear whether the seemingly lower rates of miscarriages reported in humans is due to limited epidemiologic data leading to reporting bias or biological resistance mechanisms impeding human fetal infection. Although both are eutherian taxa utilizing placental membranes for fetal development, each has significant differences in cellular composition, blood-flow patterns, and tissue structures within the placenta. Ex vivo systems using placenta explants have provided important information regarding the pathologies and immune responses associated with other congenital infections (23, 24). Here, we use placenta explants to compare permissivity of the maternal-fetal interface of sheep and humans to infection with RVFV. Furthermore, we use the explants to screen the replicative capacity of LAVs, given their variable abilities to cause abortions and teratogenic effects in livestock in vivo (10, 11, 16-19). Since limited epidemiological data exists to fully understand the impact of vertical transmission of RVFV in pregnant women, ex vivo analyses can play a pivotal role in risk assessment.

## Results

To provide a relevant comparison between human and sheep placenta with regards to permissivity of semi-intact structures located at the maternal-fetal interface, it is important to delineate the structural differences between placenta of humans and sheep. Unlike human placentas, which are discoidal, sheep placentas have a cotyledonary structure containing up to 70-100 individual fetal-derived cotyledons that interdigitate with maternal caruncle microvilli constituting what are known as placentomes (**Figure 1A**). Villi, which are tree-like branches made of trophoblasts and fetal vasculature, play an important role in nutrient and oxygen exchange between the fetus and mother. The placentomes are distributed across the entire gestational sac and connected by the allantoic and chorionic membranes (25). The fetal membranes provide a protective barrier between the fetus and mother. Each placentome fits inside a specialized pocket within the maternal decidua, called a caruncle. The trophoblast-lined cotyledon connects with the decidua through weak invasion by cytotrophoblasts. The cytotrophoblasts fuse with, or invade into, the uterine epithelial cells, creating an epitheliochorial barrier that defines livestock placenta.

**Figure 1.**
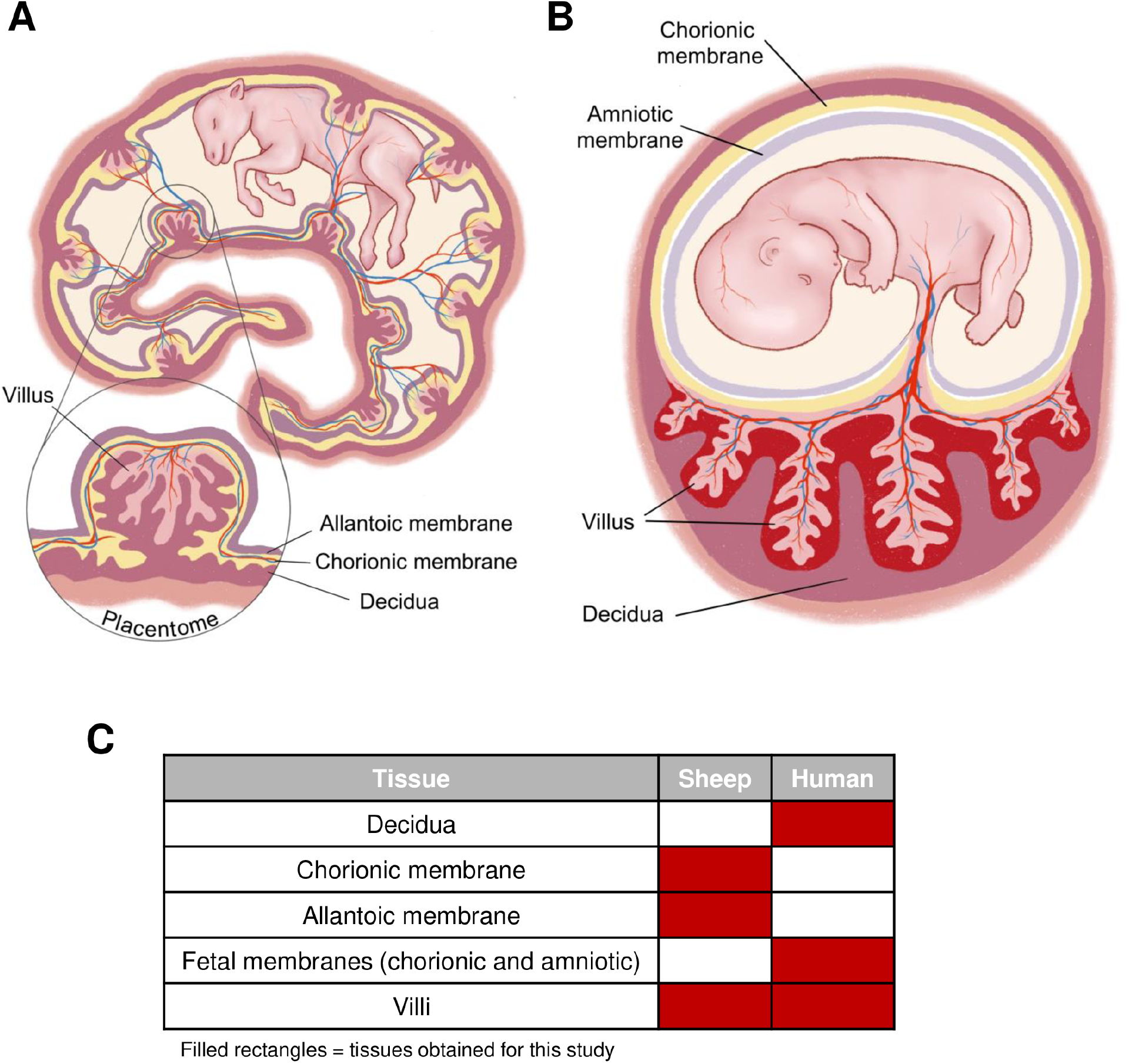
Sheep and human placental structure. A) Diagram depicting the structures that make up the sheep placenta. Sheep contain 70-100 placentomes that cover the gestational sac. The placentome consists of the villous, allantoic membrane, chorionic membrane, and decidua. B) Diagram depicting the structures that make up the human placenta. The human placenta consists of the fetal-derived villous and allantoic membrane, and chorionic membrane (fetal membranes) which is attached to the maternal decidua. C) Table representing the sheep and human tissues collected for these studies.

Humans, in contrast, have a single, discoid placenta (**Figure 1B**). Like sheep, villi of the human placenta contain fetal vasculature and are lined by trophoblast. Unlike sheep, trophoblast lining human placental villi are bathed in flowing maternal blood which is critical to provide nutrients and oxygen to the fetus. The chorioamniotic membrane surrounds the developing fetus, maintains the amniotic fluid, protects the fetus from injury, and regulates fetal temperature. Decidualized endometrium at the implantation site, known as the decidua basalis, comes in direct contact with villi via extravillous trophoblasts that invade the endomyometrium to establish vascular flow to the developing placenta. Blood flows from adapted uterine vessels in the decidua into the intervillous space.

Separation and dissection of livestock placenta into villi, chorionic membrane, and allantoic membrane (26-28) and human placenta into villi, fetal membranes and decidua is feasible (23, 29) **(Figures 1C**). Maternal decidual tissues are generally not available from sheep after natural birth and thus are not analyzed in this study. We hypothesize that culturing placental tissue sub-types will allow identification of regions of the maternal-fetal interface that are susceptible to RVFV infection.

### Sheep placenta explants are permissive to infection with RVFV vaccine strains

Term placentas and one preterm placenta were collected from four sheep housed at a local farm in western Pennsylvania (**Table 1**). Dissected villi, chorionic membranes, and allantoic membranes were inoculated with 1 x 10^5^ plaque forming units (pfu) of wild-type RVFV (ZH501) or attenuated strains (ΔNSm, ΔNSs, MP-12, or ΔNSs/ΔNSm) (**Figure 2A-B**). Culture supernatant was collected every 12-24 hours to generate a viral replication curve by q-RT-PCR (**Figure 2C**).

**Table 1:** Sheep placenta demographics.

| Sheep | Breed | Gestation | Offspring Sex |
| --- | --- | --- | --- |
| Donor 1 | Texel x Suffolk | Approx. 4mo | - |
| Donor 2* | Texel x Suffolk | Term | Two males |
| Donor 3 | Texel x Dorset | Term | Male |
| Donor 4* | Texel x Suffolk | Term | Two males |

x crossed breed, \* twins, - unknown

**Figure 2.**
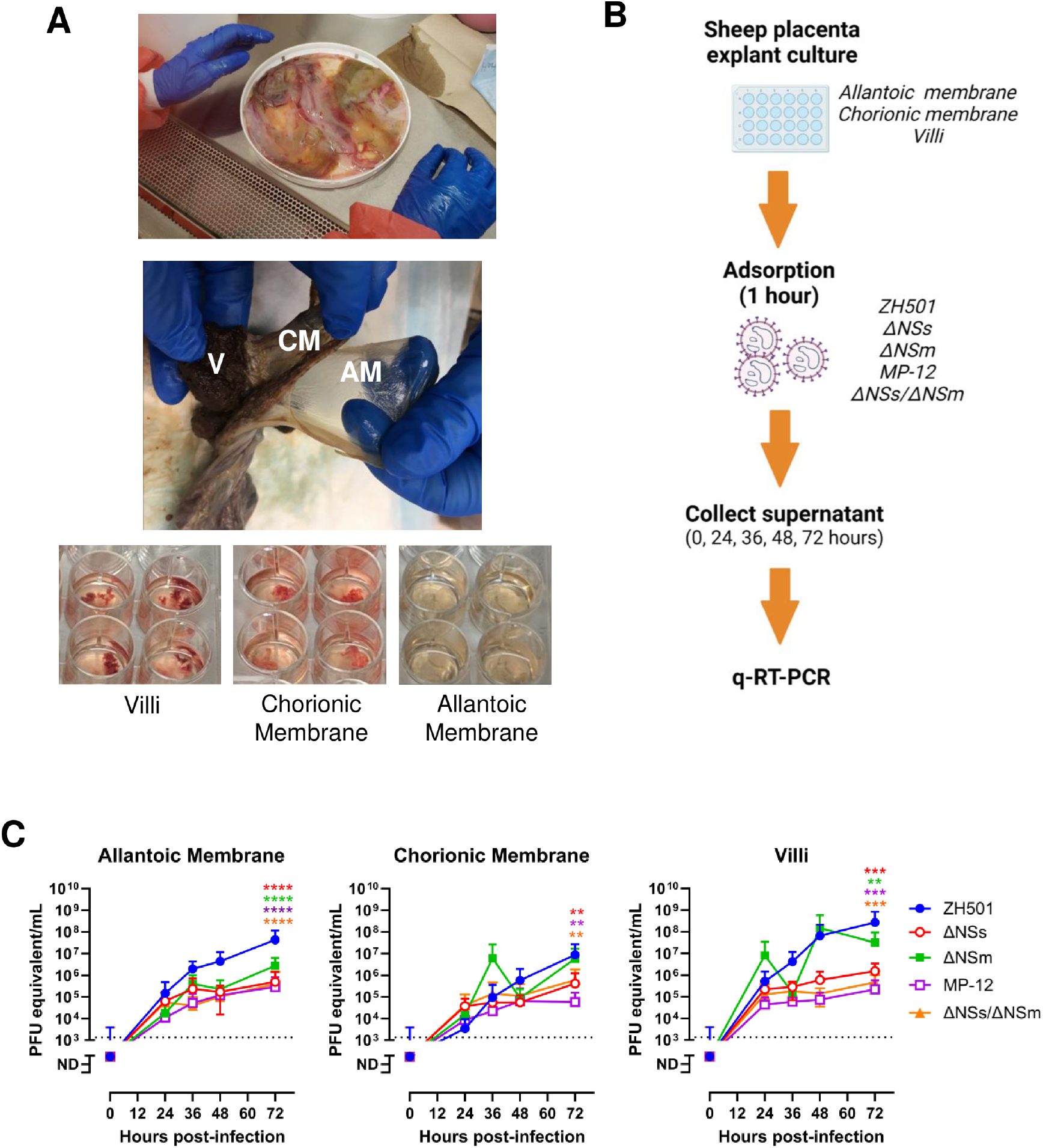
Sheep placenta explants are permissive to infection with RVFV. A) Representative image of a whole (top), separated (middle; formalin fixed), and dissected (bottom) sheep placenta. 5 x 5 mm sections of the villi (bottom, left), chorionic membrane (bottom, middle), and allantoic membrane (bottom, right) were cultured in 24-well plates. B) Diagram depicting the timeline for sheep placenta explant culture infections. Tissue dissections were inoculated with 1 x 10^5^ pfu RVFV ZH501, ΔNSs, ΔNSm, MP-12, ΔNSs/ΔNSm for 1 hour (n = 12 each). Virus was removed and washed prior to the addition of culture growth medium. Culture supernatant was collected at 0, 24, 36, 48, and 72 hpi. C) q-RT-qPCR was performed to quantitate virus production over time. Dashed line = limit of detection (LOD). Statistical significance was determined by a one-way ANOVA compared to ZH501 cultures. * p<0.05, ** p<0.01, *** p<0.001. **** p<0.0001. n.s. = not significant. Red, green, purple, and orange asterisks designate the p-value of ΔNSs, ΔNSm, MP-12, ΔNSs/ΔNSm compared to ZH501, respectively.

Villi and both fetal membranes obtained from sheep were permissive to infection with all strains of RVFV (**Figure 2C**). Viral RNA was detected as early as 24 hours post-infection (hpi) and increased through 72 hpi. RVFV ZH501 and ΔNSm produced the highest levels of viral RNA in the culture supernatant compared to the other strains, with some cultures reaching as high as 5 x 10^8^ pfu equivalents/mL (**Figure 2C**). For wild-type ZH501, there was a steady 3-log increase in virus over 72 hours for all tissues, whereas the replication curve for the villous and chorionic membrane cultures infected with ΔNSm showed a biphasic response. After infection, villi consistently shed more RVFV ZH501 and ΔNSm compared to the allantoic and chorionic membranes. The strains with the lowest tissue replication in sheep explants were ΔNSs, MP-12 and ΔNSs/ΔNSm (approx. 1 x 10^4^ -1 x 10^6^ pfu equivalents/mL) and replication appeared to plateau after 24 hours post-infection (**Figure 2C & Supplemental Figure 1)**.

Consistent with the q-RT-PCR results, immunohistochemistry revealed widespread infection within the cytoplasm of cells throughout the villi and allantoic and chorionic membranes inoculated with ZH501 as depicted by moderate levels of RVFV nucleoprotein immunopositivity (**Figure 3 & Supplemental Table 1**). Immune cells with detectable viral nucleoprotein were also observed in the stroma of the tissues infected with ZH501. For ΔNSm, strong nucleoprotein immunopositivity was present in the allantoic membrane and villous in addition to stromal immune cells. The presence of nucleoprotein in the chorionic membrane was inconclusive due to artifact of background reactivity. For the other attenuated RVFV strains, moderate levels of cytoplasmic and membranous viral antigen was detected in the chorionic membrane inoculated with MP-12, and immune cells containing nucleoprotein were rarely identified in the stroma and trophoblasts. Low levels of immunopositivity was observed in the chorionic membrane and none was detected in the allantoic membrane of tissues exposed to ΔNSs/ΔNSm in culture. In contrast, strong nucleoprotein deposition was seen in the cytoplasm of cells within the villous, and RVFV antigen-positive immune cells were also observed in the stroma of the villous of ΔNSs/ΔNSm infected tissues. ΔNSs, however, did not appear to infect sheep placenta explant cultures as negligible antigen staining was identified. Overall, ZH501 and ΔNSm caused widespread infection of the fetal membranes and villous. MP-12 and ΔNSs/ΔNSm infect the chorionic membrane and/or villous to varying degrees.

**Figure 3.**
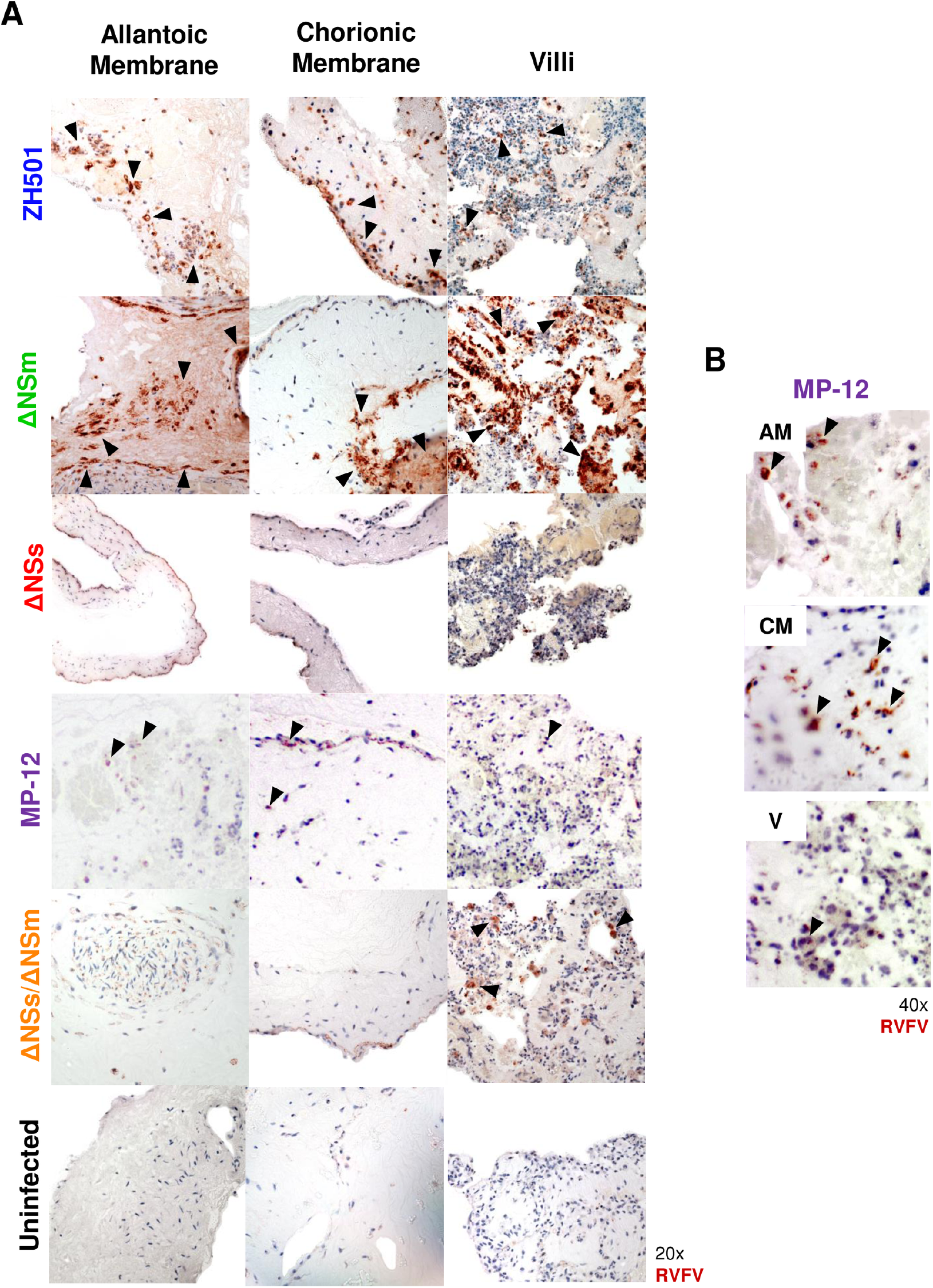
ZH501 and ΔNSm undergo wide-spread infection in sheep placenta explants. Immunohistochemistry (IHC) images of sheep allantoic membrane (AM), chorionic membrane (CM), and villi (V) cultures infected with RVFV ZH501, ΔNSm, ΔNSs, MP-12, ΔNSs/ΔNSm or uninfected controls. A) Images were taken at 20x magnification (left) by light (optical) microscopy. B) 40x magnification images of sheep placenta cultures infected with MP-12 (right). Red-brown = RVFV nucleoprotein. Blue staining = cell structural counterstain. Black arrowheads highlight regions infected with RVFV.

### All vaccine strains infect human villous explants, but tissue distribution differs in viruses lacking the viral NSs protein

Human placentas from livebirths were collected from women near-term (32- and 36-week gestation) (**Table 2**). Dissected decidua, fetal membranes, and villi were inoculated with 1 x 10^5^ pfu of RVFV ZH501, ΔNSm, ΔNSs, MP-12, or ΔNSs/ΔNsm and culture supernatant was collected over time to generate a viral replication curve (**Figure 4A-B**).

**Table 2:** Human placenta demographics.

| Human | Gestation | Offspring Sex | Notes |
| --- | --- | --- | --- |
| Donor 1 | 35 weeks, 6 days | Female | Peripheral biopsy of placenta accreta specimen |
| Donor 2* | 32 weeks, 4 days | Female | Dichorionic, diamniotic twins |
| Donor 3* |  | Male | Suspected intrauterine growth restriction (male) |

\* twins

**Figure 4.**
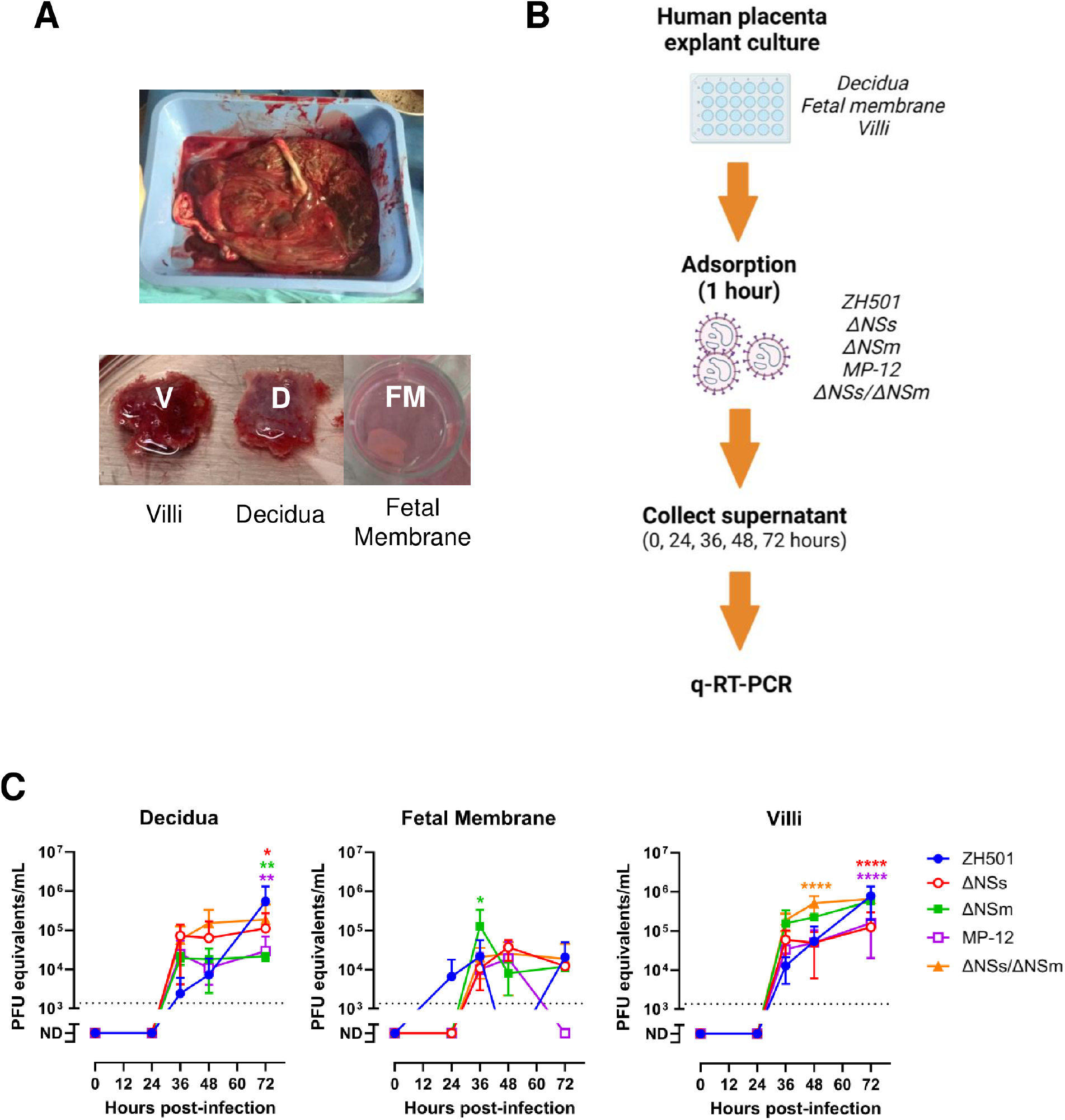
Human placenta explant cultures are permissive to RVFV strains. A) Representative image of a whole (top) and dissected (bottom) human placenta and decidua. B) Experimental timeline for human placenta explant culture infections. Tissue dissections were inoculated with 1 x 10^5^ pfu RVFV ZH501, ΔNSs, ΔNSm, MP-12, ΔNSs/ΔNSm for 1 hour (n = 3-5 each). Virus was removed and washed prior to the addition of culture growth medium. Culture supernatant was collected at 0, 24, 36, 48, and 72 hpi. C) q-RT-PCR was performed to quantitate virus production over time. Dashed line = limit of detection (LOD). Statistical significance was determined by a one-way ANOVA compared to ZH501 cultures. * p<0.05, ** p<0.01, *** p<0.001. **** p<0.0001. n.s. = not significant. Red, green, purple, and orange asterisks designate the p-value of ΔNSs, ΔNSm, MP-12, ΔNSs/ΔNSm compared to ZH501, respectively.

Like the sheep cultures, ZH501 and ΔNSm generated the highest levels of viral RNA over time, with some human explant cultures reaching 10^6^ pfu equivalents/mL. ΔNSs and MP-12 reached just above 10^5^ pfu equivalents/mL in villous cultures by 72 hours (**Figure 4C & Supplemental Figure 2**). Surprisingly, villous and decidua cultures with the double mutant virus, ΔNSs/ΔNSm, reached similar levels of viral RNA as the ZH501 cultures by 72 hours. Overall, virus within villi and decidua replicated at greater efficiency than the fetal membrane as demonstrated by high viral RNA. Fetal membrane cultures reached approximately 5 x 10^4^ pfu equivalents/mL at endpoint, whereas villi and decidua produced up to 7 x 10^5^ pfu equivalents/mL of viral RNA.

Virus was detected in human villous tissue inoculated with ZH501 or attenuated virus strains by immunohistochemical analysis (**Figure 5A**). Immunopositivity was observed in a linear pattern in syncytiotrophoblast and/or within villous stromal cells. Although the number of immunopositive villi appeared fewer in MP-12 and ΔNSs/ΔNSm samples compared to ZH501, additional data are needed to quantify differences in infection kinetics. Immunopositivity was not observed in uninfected villous controls. (**Supplemental Table 2**).

**Figure 5.**
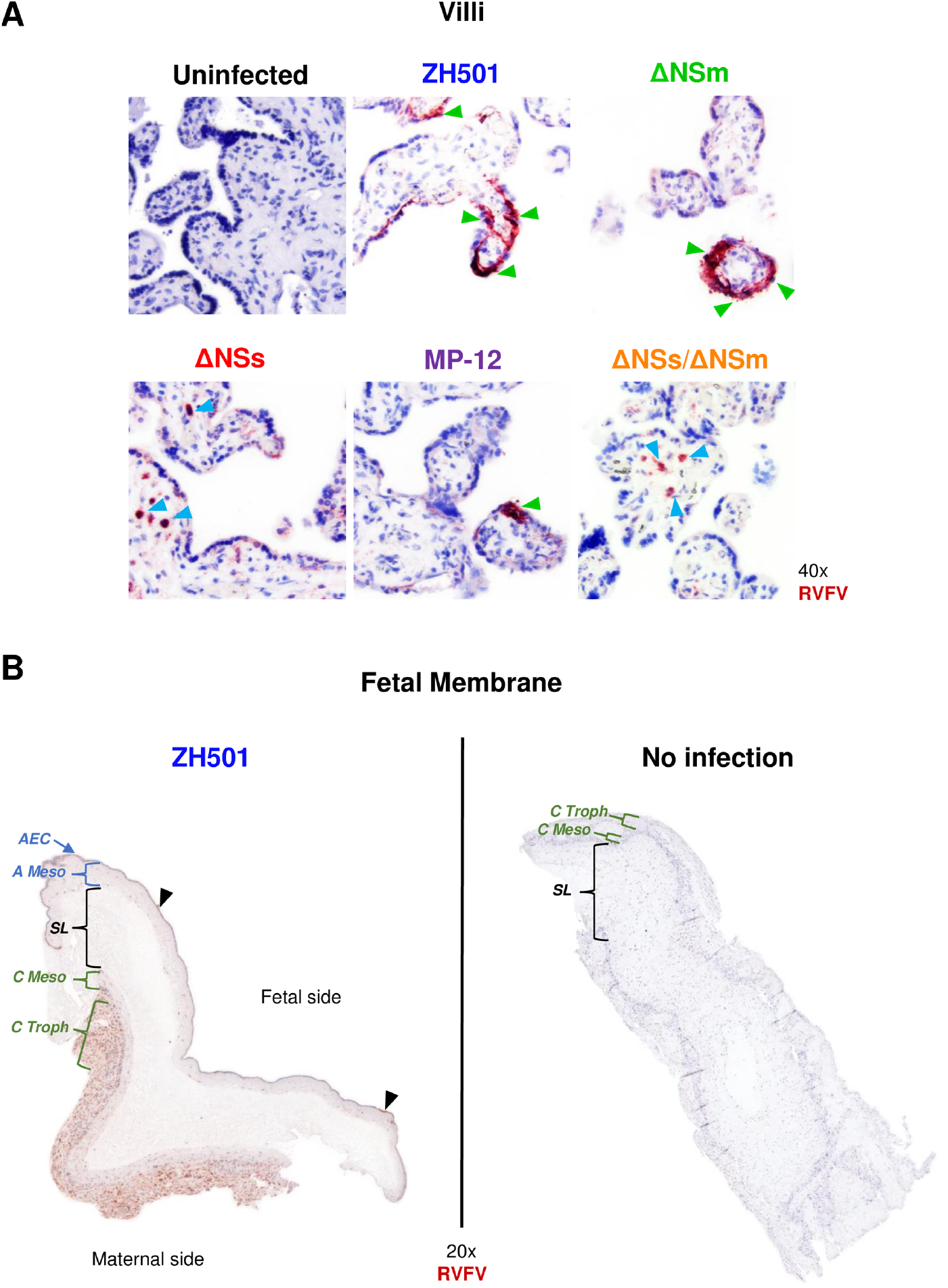
RVFV ZH501 and attenuated strains infect human placenta villous explants and RVFV ZH501 infects the maternal-facing side of the fetal membrane. A) Immunohistochemistry images of human villous cultures infected with RVFV ZH501, ΔNSm, ΔNSs, MP-12, ΔNSs/ΔNSm or uninfected controls. Images were taken at 40x magnification by light microscopy. Green arrowheads highlight regions of syncytiotrophoblasts infected with RVFV. Blue arrowheads highlight intravillous cells infected with RVFV. B) Immunohistochemistry images of ZH501 infected (left) or uninfected (right) fetal membrane cultures. Full tissue scans were performed at 20x magnification. Structures of the fetal membrane are highlighted by brackets. Structures on the fetal side include the amnion epithelial cells (AEC) and amniotic mesoderm (A Meso). The spongy layer (SL) separates the fetal and maternal sides. Structures on the maternal side include the chorionic mesoderm (C Meso) and the chorionic trophoblasts (C Tropho). Black arrowheads highlight punctate regions infected with RVFV. Red-brown = RVFV nucleoprotein. Blue staining = cell structural counterstain.

In contrast to villi, human decidua and fetal membrane cultures inoculated with ZH501 or the attenuated strains demonstrated weak and/or variable immunopositivity. One fetal membrane culture infected with wild-type ZH501 displayed moderate to strong staining in the maternal-facing layers (chorionic trophoblast and decidua capsularis; **Figure 5B & Supplemental Figure 3A**). This result may indicate regional differences in cellular susceptibility to infection across the fetal membrane.

### RVFV NSs antagonizes production of both Type-I and -III interferons in human villous cultures

Innate anti-viral responses, such as the expression of type-I and type-III interferons (IFNs), can impact viral replication and/or dictate pathogenesis (30). Type-I interferons are expressed by most cells to provide systemic control of infection, whereas type-III interferons are expressed at barrier surfaces, such as placental trophoblasts, to control infections locally (31, 32). Type-I interferons primarily provide protection during the first trimester of pregnancy in humans, whereas type-III interferons are more highly expressed and contribute to an anti-viral state during the third trimester (33). RVFV NSs is a type-I IFN antagonist, and its role at the maternal-fetal interface has not been evaluated. Furthermore, the antagonistic properties of NSs have not been evaluated in the context of type-III IFN. To determine whether there were strain-specific differences in expression of IFN-α or IFN-λ1, ELISAs were performed to quantify protein levels within end-point culture supernatant. Wild-type ZH501 did not induce IFN-α or IFN-λ1 expression in human placental samples (**Figure 6**). In contrast, for attenuated strains lacking NSs expression (ΔNSs and ΔNSs/ΔNSm), IFN-α was elevated in the supernatant of human villous and decidua cultures, with higher concentrations in the villi compared to the decidua. IFN-λ1 is elevated in the supernatant of villous cultures infected with ΔNSs, MP-12 and ΔNSs/ΔNSm compared to uninfected controls, which had undetectable levels of IFN-λ1 (**Figure 6B**). IFN-λ1 expression was relatively low or not detected in decidua and fetal membrane cultures infected with LAVs. Overall, viruses without the NSs protein induce higher levels of type-I and -III interferons in human villous explant cultures, which demonstrates that NSs is an antagonist of both type of IFN in the placenta.

**Figure 6.**
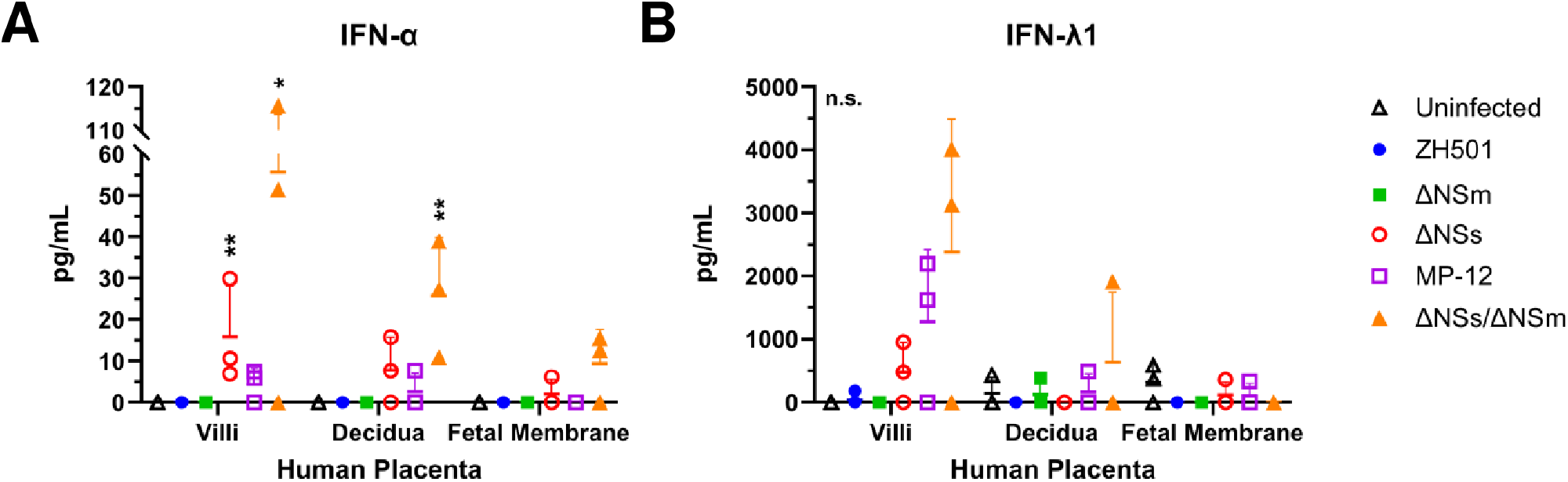
Attenuated RVFV LAVs induce Type I and III IFNs in human placenta. Protein levels of IFN-α (left) or IFN-λ1 (right) were measured in culture supernatant collected at endpoint (48 or 72 hpi) by ELISA. Culture supernatant was collected from villus, decidua, or fetal membrane cultures inoculated with RVFV ZH501, ΔNSm, ΔNSs, MP-12, ΔNSs/ΔNSm (n = 3-5). IFN-α limit of detection (LOD) = 3.2 pg/mL. IFN-λ1 LOD = 13.72 pg/mL. Statistical significance was determined by a two-way ANOVA compared to uninfected cultures. * p<0.05, ** p<0.01. n.s. = not significant.

### Sheep placenta explant cultures produce more virus than human placenta explant cultures

Overall, sheep placenta cultures produced more virus (2 x 10^5^ – 5 x 10^8^ pfu equivalents/mL) than human cultures (1 x 10^4^ – 7 x 10^5^ pfu equivalents/mL) (**Figure 2C & 4C**). To evaluate the replication kinetics across viruses and tissues, we calculated the slope of the increase in viral RNA in each culture type by linear regression (**Figure 7**). We started at 24 hpi for sheep samples or 36 hpi for human samples due to the delay in replication kinetics (**Figures 2C, 4C**). The decidua was only collected from human placentas, and thus there is not a sheep equivalent for comparison. However, in the human decidua, wild-type ZH501 clearly had the steepest slope (i.e. highest virus replication rate), whereas the LAVs did not produce virus after 36 hpi, as portrayed by the near zero or slightly negative slope (**Figure 7, left panel**). For the fetal membranes (**Figure 7, middle panel**), sheep cultures had steeper slopes compared to human cultures across all virus strains. All LAVs had a positive slope in sheep allantoic membrane cultures, whereas ΔNSs and MP-12 did not grow well after 36 hpi (slope: -0.007 and -0.111, respectively) in chorionic membranes. ΔNSs/ΔNSm grew quite well in sheep chorionic membranes. For all strains, including ZH501, the virus replication rates were near zero or negative within human fetal membrane cultures. Villi were highly permissive to virus replication for all strains in both sheep and human samples (**Figure 7, right panel**), except for ΔNSs, which did not grow in human villous cultures. Taken together, using the slope of the increase in viral RNA over time provides insight into the relative permissivity of sheep and human maternal-fetal interface structures.

**Figure 7.**
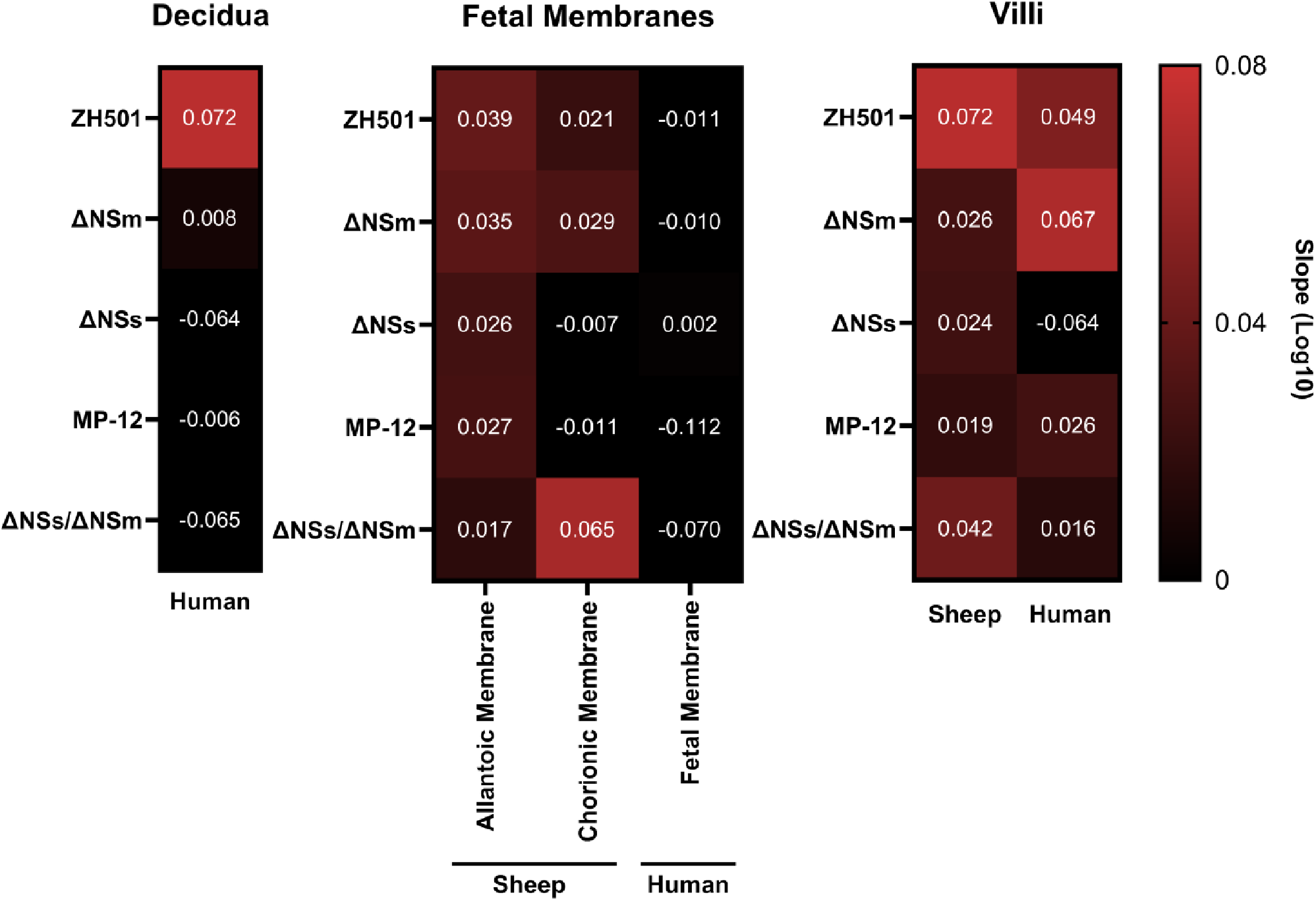
The rate of virus production is similar between hosts, despite delayed production in human cultures. A simple linear regression was performed to calculate the rate of virus production (slope) from virus growth curves for decidua, fetal membrane, and villous cultures starting at 24 hpi (sheep) and 36 hpi (humans). Positive slope = red shades; no slope = black. Individual slopes are depicted by white numbers.

## Discussion

There are multiple mechanisms by which pathogens can undergo vertical transmission, an event in which a fetus is infected through exposure from its infected or colonized mother. TORCH pathogens (*Toxoplasma gondii*, other, rubella virus, cytomegalovirus, herpes simplex virus) can adversely affect the developing human fetus through several potential mechanisms including transplacental transmission, direct insult to the placenta, fetal-maternal hemorrhage, transmission across fetal membranes, or ascending sexual transmission. Pathogens also induce inflammatory responses at the maternal-fetal interface that adversely affect the fetus (31, 34). The major structures at the maternal-fetal interface that protect against vertical transmission are the maternal decidua and fetal-derived membranes (amnion and chorioallantois) and villi (**Figure 1**). For many pathogens, the mechanism of vertical transmission across the maternal and fetal structures is incompletely understood. Although RVFV has caused massive abortion storms in livestock since the year 1930 (35, 36), few studies have examined the cellular tropism at the maternal-fetal interface or the routes that RVFV could take to reach the fetus in utero.

In a previous study, Odendaal et al. examined placentas from naturally aborted sheep during a 2010-2011 RVF outbreak in South Africa (37). Extensive RVFV antigen immunostaining was found within trophoblasts in the cotyledonary chorioallantois. In the intercotyledonary areas, such as the chorioallantoic membrane, RVFV antigen was sparse yet detectable. Significant necrosis was observed in the trophoblasts and endothelial cells of the cotyledonary and intercotyledonary chorioallantois, whereas necrosis of the villous trophoblasts was typically diffuse. In a separate experimental study in pregnant sheep, inoculation with the wild-type RVFV strain 35/74 at one-third gestation resulted in abortion at 6 dpi, and immunostaining showed wide-spread infection of maternal epithelial cells and fetal trophoblasts lining the hemotophagous zone of the placenta (2). Placentas showed signs of hemorrhage of the maternal villi, extensive necrosis of the maternal epithelium, and necrosis of the fetal trophoblasts; analyses of the fetal membranes were not performed in this study. These two studies provide important insights into RVFV infection of the maternal-fetal interface of livestock. However, experimental studies in pregnant sheep are expensive, cumbersome, and require levels of agricultural animal biocontainment that are restrictive. Thus, given the limited data from both natural and experimental RVFV infection during pregnancy in sheep, we sought to develop tractable placental explant cultures for direct comparison with their human counterparts.

There is increasing clinical evidence of vertical transmission of RVFV during human pregnancy and epidemiologic data demonstrating a clear association with poor pregnancy outcomes (6, 7). However, histopathological studies of human fetuses and placenta were not performed in these studies. To overcome this limitation, we previously inoculated human second trimester placenta explants with wild-type RVFV and found that villi were susceptible to infection with RVFV strain ZH501 (29). The villous cytotrophoblasts and syncytiotrophoblast were particularly permissive to RVFV. This finding was alarming because syncytiotrophoblast are generally resistant to most TORCH pathogens (38) by providing a large physical barrier (39-41) or secreting antimicrobial factors (23, 42). In a separate study, human term villous explants were also permissive to infection with RVFV strain 35/74 (2).

Given the data from our lab and others that RVFV can replicate in human explants, here we conducted a direct comparison between human and sheep tissues regarding permissivity to infection with RVFV and LAVs. We obtained villi, chorioallantoic, and allantoic membranes from freshly harvested sheep placentomes and dissected them into explants of comparable size. Sheep explant cultures containing villi produced substantial levels of RVFV ZH501 viral RNA over 72 hours – approximately 100-fold higher than that produced by human villi. All 3 of the human tissues we tested (decidua, fetal membrane, and villi) were also permissive to wild-type RVFV ZH501, although virus production had delayed kinetics and lower overall production compared to ovine counterparts. In our model of ex vivo placental infection, human villi appeared more permissive to ZH501 infection than the decidua or fetal membranes. Overall, sheep explants were more permissive to RVFV infection, and this is reflective of the very high rates of fetal infection in pregnant sheep naturally infected with RVFV. This difference in permissivity and replicative capacity could be due to differences in viral receptor expression and/or innate immune responses and warrant further mechanistic studies.

The high rates of mortality in livestock and morbidity in humans during RVFV outbreaks cause significant socioeconomic strain in affected areas (1). To reduce the impacts of RVF disease in livestock and incidental transmission to humans, live-attenuated vaccines (LAVs) have been developed for use in ruminant livestock. While LAVs are effective at preventing infection after challenge, the safety of some LAVs may be questionable as several have been shown to display residual virulence in pregnant sheep (10, 11, 16, 43). Here we used the placental explant model to study the permissivity of sheep and human tissues to infection with several relevant LAVs. In sheep placenta explants, LAVs (ΔNSs, MP-12, ΔNSs/ΔNSm) replicated to moderate levels in villi and fetal membranes, albeit significantly less than wild-type strain ZH501. ΔNSm, on the other hand, generally replicated as well as ZH501 in both sheep and human explants, which is consistent with other *in vivo* studies showing ΔNSm retains significant neurovirulence (44). Human villous cultures infected with ΔNSs, MP-12, and ΔNSs/ΔNSm undergo early replication which plateaued by 36 hpi. Consistent with these observations, villous and decidua culture supernatant infected with ΔNSs or ΔNSs/ΔNSm had elevated levels of IFN-α and IFN-λ1 72 hpi compared to uninfected and ZH501 infected controls. ZH501 and ΔNSm, which express NSs, were infectious in the villous, decidua, and fetal membrane cultures, and neither IFN-α nor IFN-λ1 were detected in the supernatant. From these results, we can conclude the nonstructural protein, NSs, contributes to virulence at the maternal-fetal interface by antagonizing the expression of type-I interferons, cytokines that contribute to an antiviral state within the cell and surrounding areas (45). These results also suggest that NSs inhibits type-III interferons in the villous which is further corroborated by another in vitro study that showed exogenous RVFV NSs inhibits interferon-stimulated response element activation in response to IFN-λ1 (46). It is promising to see human placenta explants control LAV infection; however, if these attenuated viruses are not controlled in the periphery and reach or breach the maternal-fetal interface, the local immune response could contribute to adverse effects to the developing fetus. For instance, the type-I and type-III interferon responses stimulated by the presence of RVFV might lead to adverse pathological effects as seen with Zika virus infection in mice (30).

Live-attenuated vaccines for human use have not yet been approved, but a few have advanced into clinical trials. In 2006-2008, the attenuated MP-12 strain was included in a phase I/II clinical trial, but it has yet to progress further to the authors knowledge (14). Other vaccine candidates, including the ΔNSs/ΔNSm and a 4-segmented vaccine approach (RVFV-4s), have or will soon enter early stage clinical trials in humans (21, 22). Given the evidence of vertical transmission of RVFV in pregnant women and direct infection of human placenta in culture, it is important to evaluate the risk of infection and subsequent pathogenesis of human placentas to RVFV LAVs. Furthermore, early stage vaccine clinical safety trials typically do not include pregnant women due to safety and ethical concerns, making the study of these questions in a laboratory setting even more crucial. Placenta explant cultures are also a tractable model to use in a BSL-3 setting. The use of human and sheep placenta explants, studied at various stages of gestation, could provide pivotal information regarding the cell-types targeted by RVFV and the local immune responses to infection. It is important to emphasize that these explant cultures are an incomplete yet biologically relevant model due to the lack of a circulatory system and cross-body communication. Regardless, explants provide a diverse local cellular environment, structure, and inter-tissue cellular communication that immortalized cell cultures cannot.

RVFV has caused outbreaks with high rates of abortion among sheep and other ruminant livestock for almost a century (35, 36). In contrast, the apparent impacts on pregnant women appear to be significantly lower, but limited epidemiological studies examining pregnancy outcomes in women in endemic regions may account for this perceived difference. We found here that villi from both species are highly susceptible to RVFV infection as evident by the increasing levels of virus over time and the antigen detection. Once RVFV reaches the chorionic villi, it would likely be difficult to control infection and spread to the fetus given the proximity to the fetal endothelium and blood supply (25). Interestingly, the fetal membranes from sheep produced high levels of virus, whereas RVFV did not replicate well in human fetal membrane cultures. Looking closer at one of the human fetal membranes, there is a stark difference in the regions that are infected with RVFV ZH501. The chorionic trophoblasts, which naturally face the mother, were diffusely infected with ZH501, whereas the amniotic epithelium, which faces the fetus, had very little infection. The nature of the explant culture provides equal exposure of all sides of the explant to virus, preventing an infection bias. The fetal membrane in humans, particularly the amniotic epithelium and stroma, which only had a few regions of punctate infection, may play an important role in preventing vertical transmission and protecting the fetus. Surprisingly, fetal membrane cultures did not produce high concentrations of IFN-α or IFN-λ1, suggesting the resistance to infection could be mediated by a different mechanism. Sheep allantoic and chorionic membranes appear to be equally vulnerable to infection which might explain the higher incidence of fetal demise in livestock compared to humans. *In vivo* studies using reporter viruses in pregnant sheep and rats could delineate the spatial and temporal aspects of vertical transmission of RVFV. This study is limited by the low number of biological replicates due to difficulty in procuring sheep and human placentas, thus additional studies are warranted to corroborate our findings. Furthermore, given multiple studies have shown that virus dose and gestational age may influence disease outcome (2, 10, 16), future studies should be performed to determine whether these factors could affect vertical transmission across the placenta. Further explant studies should also be performed to understand what makes human amniotic epithelial cells more resistant to RVFV in order to develop targeted therapeutics to prevent or minimize the pathogenic effects following vertical transmission.

In summary, we have used sheep and human placenta explant cultures as a tool to identify cells and tissues targeted by RVFV at the maternal-fetal interface. We provide evidence that although LAVs can infect the decidua, fetal membrane, and villi of humans, spatial restraints, particularly the decidua and fetal membrane, could play an important role in controlling infection and preventing virus dissemination to the fetus.

## Materials and methods

### Viruses

Virulent RVFV strain ZH501 was generated from reverse genetics plasmids. The recovered virus was provided to our laboratory by B. Miller (CDC, Ft. Collins, CO) and S. Nichol (CDC, Atlanta, GA). RVFV strains with gene deletions (ΔNSs, ΔNSm, ΔNSs/ΔNSm) were provided by A.K. McElroy (University of Pittsburgh). MP-12 was generously provided by Ted Ross (Cleveland Clinic Florida Research & Innovation Center, Port St. Lucie, FL). All strains of RVFV were propagated on Vero E6 cells following standard methods and the stock titer was determined by standard viral plaque assay explained in McMillen *et al*. (29) For explant studies, stock virus was diluted in virus growth medium (DMEM, 2% (v/v) FBS, 1% L-glutamine, and 1% penicillin-streptomycin) to the desired dose. Uninfected controls were mock inoculated with virus growth medium.

### Biosafety

Research with RVFV strains ZH501, ΔNSs, and ΔNSm was performed at biosafety level 3 (BSL-3) in the University of Pittsburgh Regional Biocontainment Laboratory (RBL). The University of Pittsburgh RBL is registered with the Centers for Disease Control and Prevention and the United States Department of Agriculture for work with select agents. Work with RVFV strains MP-12 and ΔNSs/ΔNSm which are both excluding from the DSAT Select Agent lists was performed at biosafety level 2 (BSL-2) as per local Institutional Biosafety Committee (IBC) guidelines.

### Placenta procurement

Sheep (Texel crossed with Dorset or Suffolk breeds) placentas were collected by George and Lisa Wherry at Wherry Farms, Scenery Hill, PA. Placentas were collected from the field or barn, or extracted directly from the sheep. Most of the sheep placentas were procured at term (n = 3; approximately 5 months gestation) after a natural delivery, whereas one placenta was obtained from a pre-term delivery (4 months gestation). Immediately after collection, placentas were placed in complete growth media consisting of DMEM/F12 (1:1), 10% FBS, 2% pen-strep, and 250 μg/mL of amphotericin B to minimize the growth of microbial contaminants. Human placentas isolated between 32-36 weeks gestation from C-section were obtained from the Steven C Caritis Magee Obstetric Maternal and Infant Biobank through an honest broker system within 30 minutes of delivery.

### Ethics Statement

Human tissue procurement was approved by the University of Pittsburgh Institutional Review Board (IRB) 19100322.

### Explant studies

All placentas were dissected within 6-12 hours of procurement, and dissection was conducted based on tissue type (humans: decidua, villi, fetal membranes (chorion and/or amnion); sheep: chorionic membrane, allantoic membrane, villi). Placenta tissue sections, approximately 5 x 5 mm sections, were placed in 24-well plates, covered in complete growth medium and allowed to rest overnight at 37°C, 5% CO_2_. On the day of infection, the complete growth media was removed from each placenta section prior to inoculation with 200 μL of RVFV (1 x 10^5^ pfu).

Placenta sections were incubated for 1 hour at 37°C, 5% CO_2_ to allow for virus adsorption. Following the adsorption period, the inoculum was removed, tissue were washed once with 1x PBS, and then 1 mL of virus growth medium was added to all samples. The plate was cultured for 48 or 72 hours and 100 μL of culture supernatant was pooled from 1-2 wells from each tissue daily at 0, 24, 36, 48, and 72 hpi (29). For each human placenta donor (n=3) 1-2 samples were analyzed from pooled culture supernatant at each time point. For each sheep placenta donor (n=4) three samples were analyzed from pooled culture supernatant at each time point.

### RNA isolation and q-RT-PCR

The culture supernatant (50 μL) was inactivated in 900 μL of TRIzol reagent (Invitrogen). RNA isolation and q-RT-PCR were performed as described (29).

### Enzyme-linked immunosorbent assay (ELISA)

Supernatant collected from human tissues at 48 or 72 hpi after infection with RVFV ZH501 or LAVs (ΔNSs, ΔNSm, MP-12, ΔNSs/ΔNSm) was used to detect total human interferon alpha (IFNα; Invitrogen catalog #BMS216) and human interleukin 29 (IL-29/IFN-λ1; abcam # ab100568) following the manufacturer’s instructions. Standard curves were generated to quantitate protein concentrations. The limit of dection (LOD) for the IFN-α and IFN-λ1 assays were 3.2 pg/mL and 13.72 pg/mL, respectively. For each human placenta donor (n=3) 1-2 samples were analyzed from pooled culture supernatant at either time point.

### Immunohistochemistry and microscopy

Human and sheep tissues were fixed in 4% PFA for 24 hours then paraffin embedded by standard protocols. Tissue sections 6 μm thick were embedded on slides and baked overnight at 65°C. Slides were deparaffinized using a standard xylenes and alcohol rehydration series, then boiled in 10 mM citric acid buffer (pH 6.0) to unmask antigen-binding epitopes. Tissue sections were then permeabilized using 0.1% Triton X-100 detergent in PBS. Tissue sections were blocked in BLOXALL blocking solution (Vector Laboratories) for 10 minutes, then washed 3x in PBS. Then using the VECTASTAIN Elite ABC Kit, Peroxidase (Rabbit IgG) and Vector NovaRED Substrate Kit, Peroxidase (Vector Laboratories), tissue sections were blocked in normal blocking serum, avidin block, and biotin block for 15-20 minutes, with five washes in 1x PBS between each block. Following five washes in PBS, anti-RVFV nucleoprotein rabbit IgG (1:100 dilution; custom made via Genscript) was applied to each sample and incubated for 1 hour. Tissues were washed five times in PBS, then the Vector Biotinylated Goat anti-Rabbit secondary antibody was applied for 30 minutes. Tissues were washed five times with PBS then the VECTASTAIN ABC-HRP reagent was added for 30 minutes. The tissues were washed five times, then the Vector NovaRED was applied for 15 minutes. The Vector NovaRED was removed by dunking slides into fresh water. Tissues were counterstained in 50% diluted hematoxylin (Leica, catalog # 3801575) for 20 seconds, washed in water, dunked in 5% acetic acid five times, washed in water, then incubated in Scott’s tap water (Cancer Diagnostics Inc., catalog #CM4951W) for 10 minutes to promote counterstain bluing. After dehydrating in Clear-Rite (Thermo Scientific, catalog #6901) coverslips were mounted over the tissues using toluene. Slides dried overnight before imaging. All steps were performed at room temperature unless stated otherwise. Primary delete and no infection controls were used to identify non-specific binding of the secondary detection antibody or all reagents, respectively. Red immunopositivity observed above the no infection control was deemed as positive for RVFV antigen. Hematoxylin staining provided a structural stain to identify immune cells. IHC slides were imaged by light (optical) microscopy using a Nikon NI-E confocal microscope at the University of Pittsburgh Center for Biological Imaging or the Olympus LCmicro Software at the One Health Institute at the University of California Davis. Denoising and contrasting were performed using Adobe Photoshop. Tissue samples were blinded and analyzed by a veterinary pathologist or a licensed pathologist specializing in placental pathology.

### Statistics

For virus replication curves and cytokine expression assays, one-way or two-way ANOVA with multiple comparisons were performed using GraphPad Prism (version 8.0). To calculate virus replication rates, a simple-linear regression from log(10) transformed data was performed starting at 24 or 36 hpi for sheep and human cultures, respectively.

## Supporting information

All Supplemental figures

## Acknowledgements

The authors would like to thank the Center for Biological Imaging, particularly Michael J Calderon and Katherine E. Helfrich, for assistance with image acquisition. We recognize the McGowan Center for Regenerative Medicine for histology support. Hayley Nordstrom provided exceptional original artwork. This work would not have been possible without the generous help from George and Lisa Wherry at Wherry Farms, Scenery Hill, PA. All data needed to evaluate the conclusions in the paper are present in the paper and/or the Supplementary Materials. Additional data related to this paper may be requested from the authors. B.H.B is an inventor of patents describing the development of the ΔNSs-ΔNSm-rZH501 vaccine candidate technology (USPTO: 8,673,629, 9,439,935, 10,064,933). The remaining authors declare that they have no competing interests. All work was supported by the National Institutes of Health (R01 AI150792 and R01 AI150792S1 to A.L.H., T32 AI060525 and K01 AI165965 to C.M.M). NIH award (UC7AI180311) from the National Institute of Allergy and Infectious Diseases (NIAID) supported the Operations of The University of Pittsburgh Regional Biocontainment Laboratory (RBL) within the Center for Vaccine Research (CVR). The funders had no role in study design, data collection and analysis, decision to publish, or preparation of the manuscript. Conceptualization: Cynthia M. McMillen, Amy L. Hartman; Data curation: Cynthia M. McMillen, Ryan M. Hoehl, Devin A Boyles, Jackson J. McGaughey, Lauren Skvarca, Amy L. Hartman; Formal analysis: Cynthia M. McMillen, Christina Megli, Rebecca Radisic, Lauren B. Skvarca, Brian H Bird, Amy L. Hartman; Funding acquisition: Amy L. Hartman; Investigation: Cynthia M. McMillen, Rebecca Radisic, Lauren B. Skvarca, Ryan M. Hoehl, Devin A. Boyles, Jackson J. McGaughey, Amy L. Hartman; Methodology: Cynthia M. McMillen, Christina Megli, Rebecca Radisic, Lauren B. Skvarca, Ryan M. Hoehl, Jackson J. McGaughey, Brian H. Bird, Anita K McElroy; Project administration: Amy L. Hartman; Supervision: Amy L. Hartman; Validation: Amy L. Hartman; Visualization: Cynthia M. McMillen, Rebecca Radisic, Lauren B. Skvarca, Brian H. Bird, Amy L. Hartman; Writing – original draft: Cynthia M. McMillen, Amy L. Hartman.

## References

1. Chengula AA, Mdegela RH, Kasanga CJ. 2013. Socio-economic impact of Rift Valley fever to pastoralists and agro pastoralists in Arusha, Manyara and Morogoro regions in Tanzania. Springerplus 2:549.

2. Oymans J, Wichgers Schreur PJ, van Keulen L, Kant J, Kortekaas J. 2020. Rift Valley fever virus targets the maternal-foetal interface in ovine and human placentas. PLoS Negl Trop Dis 14:e0007898.

3. Coetzer JA. 1977. The pathology of Rift Valley fever. I. Lesions occurring in natural cases in new-born lambs. Onderstepoort J Vet Res 44:205–11.

4. Coetzer JA. 1982. The pathology of Rift Valley fever. II. Lesions occurring in field cases in adult cattle, calves and aborted foetuses. Onderstepoort J Vet Res 49:11–7.

5. Baudin M, Jumaa AM, Jomma HJ, Karsany MS, Bucht G, Naslund J, Ahlm C, Evander M, Mohamed N. 2016. Association of Rift Valley fever virus infection with miscarriage in Sudanese women: a cross-sectional study. Lancet Glob Health 4:e864–e871.

6. Adam I, Karsany MS. 2008. Case report: Rift Valley Fever with vertical transmission in a pregnant Sudanese woman. J Med Virol 80:929.

7. Arishi HM, Aqeel AY, Al Hazmi MM. 2006. Vertical transmission of fatal Rift Valley fever in a newborn. Ann Trop Paediatr 26:251–3.

8. Smithburn KC. 1949. Rift Valley fever; the neurotropic adaptation of the virus and the experimental use of this modified virus as a vaccine. Br J Exp Pathol 30:1–16.

9. Muller R, Saluzzo JF, Lopez N, Dreier T, Turell M, Smith J, Bouloy M. 1995. Characterization of clone 13, a naturally attenuated avirulent isolate of Rift Valley fever virus, which is altered in the small segment. Am J Trop Med Hyg 53:405–11.

10. Makoschey B, van Kilsdonk E, Hubers WR, Vrijenhoek MP, Smit M, Wichgers Schreur PJ, Kortekaas J, Moulin V. 2016. Rift Valley Fever Vaccine Virus Clone 13 Is Able to Cross the Ovine Placental Barrier Associated with Foetal Infections, Malformations, and Stillbirths. PLoS Negl Trop Dis 10:e0004550.

11. Botros B, Omar A, Elian K, Mohamed G, Soliman A, Salib A, Salman D, Saad M, Earhart K. 2006. Adverse response of non-indigenous cattle of European breeds to live attenuated Smithburn Rift Valley fever vaccine. J Med Virol 78:787–91.

12. Lokugamage N, Ikegami T. 2017. Genetic stability of Rift Valley fever virus MP-12 vaccine during serial passages in culture cells. NPJ Vaccines 2.

13. Ikegami T, Hill TE, Smith JK, Zhang L, Juelich TL, Gong B, Slack OA, Ly HJ, Lokugamage N, Freiberg AN. 2015. Rift Valley Fever Virus MP-12 Vaccine Is Fully Attenuated by a Combination of Partial Attenuations in the S, M, and L Segments. J Virol 89:7262–76.

14. Pittman PR, Norris SL, Brown ES, Ranadive MV, Schibly BA, Bettinger GE, Lokugamage N, Korman L, Morrill JC, Peters CJ. 2016. Rift Valley fever MP-12 vaccine Phase 2 clinical trial: Safety, immunogenicity, and genetic characterization of virus isolates. Vaccine 34:523–30.

15. Pittman PR, McClain D, Quinn X, Coonan KM, Mangiafico J, Makuch RS, Morrill J, Peters CJ. 2016. Safety and immunogenicity of a mutagenized, live attenuated Rift Valley fever vaccine, MP-12, in a Phase 1 dose escalation and route comparison study in humans. Vaccine 34:424–429.

16. Hunter P, Erasmus BJ, Vorster JH. 2002. Teratogenicity of a mutagenised Rift Valley fever virus (MVP 12) in sheep. Onderstepoort J Vet Res 69:95–8.

17. Morrill JC, Carpenter L, Taylor D, Ramsburg HH, Quance J, Peters CJ. 1991. Further evaluation of a mutagen-attenuated Rift Valley fever vaccine in sheep. Vaccine 9:35–41.

18. Morrill JC, Jennings GB, Caplen H, Turell MJ, Johnson AJ, Peters CJ. 1987. Pathogenicity and immunogenicity of a mutagen-attenuated Rift Valley fever virus immunogen in pregnant ewes. Am J Vet Res 48:1042–7.

19. Bird BH, Maartens LH, Campbell S, Erasmus BJ, Erickson BR, Dodd KA, Spiropoulou CF, Cannon D, Drew CP, Knust B, McElroy AK, Khristova ML, Albarino CG, Nichol ST. 2011. Rift Valley fever virus vaccine lacking the NSs and NSm genes is safe, nonteratogenic, and confers protection from viremia, pyrexia, and abortion following challenge in adult and pregnant sheep. J Virol 85:12901–9.

20. Jenkin D, Wright D, Folegatti PM, Platt A, Poulton I, Lawrie A, Tran N, Boyd A, Turner C, Gitonga JN, Karanja HK, Mugo D, Ewer KJ, Bowden TA, Gilbert SC, Charleston B, Kaleebu P, Hill AVS, Warimwe GM. 2023. Safety and immunogenicity of a ChAdOx1 vaccine against Rift Valley fever in UK adults: an open-label, non-randomised, first-in-human phase 1 clinical trial. Lancet Infect Dis 23:956–964.

21. Rogers J. 2022. Rift Valley fever vaccines to advance with new $50 million CEPI and EU funding call, on Coalition for Epidemic Preparedness Innovations. https://cepi.net/news_cepi/rift-valley-fever-vaccines-to-advance-with-new-50-million-cepi-and-eu-funding-call/. Accessed Jan 27, 2023.

22. Rogers J. 2023. CEPI Partners with University of California, Davis to advance a vaccine against potentially deadly Rift Valley fever virus into clinical trials, on Coalition for Epidemic Preparedness Innovations. https://cepi.net/news_cepi/cepi-partners-with-university-of-california-davis-to-advance-a-vaccine-against-potentially-deadly-rift-valley-fever-virus-into-clinical-trials/. Accessed January 5, 2024.

23. Corry J, Arora N, Good CA, Sadovsky Y, Coyne CB. 2017. Organotypic models of type III interferon-mediated protection from Zika virus infections at the maternal-fetal interface. Proc Natl Acad Sci U S A 114:9433–9438.

24. Fahmi A, Brügger M, Démoulins T, Zumkehr B, Oliveira Esteves BI, Bracher L, Wotzkow C, Blank F, Thiel V, Baud D, Alves MP. 2021. SARS-CoV-2 can infect and propagate in human placenta explants. Cell Rep Med 2:100456.

25. Leiser R, Kaufmann P. 1994. Placental structure: in a comparative aspect. Exp Clin Endocrinol 102:122–34.

26. Medina L, Guerrero-Muñoz J, Castillo C, Liempi A, Fernández-Moya A, Araneda S, Ortega Y, Rivas C, Maya JD, Kemmerling U. 2022. Differential microRNAs expression during ex vivo infection of canine and ovine placental explants with Trypanosoma cruzi and Toxoplasma gondii. Acta Trop 235:106651.

27. Liempi A, Castillo C, Medina L, Galanti N, Maya JD, Parraguez VH, Kemmerling U. 2020. Comparative ex vivo infection with Trypanosoma cruzi and Toxoplasma gondii of human, canine and ovine placenta: Analysis of tissue damage and infection efficiency. Parasitol Int 76:102065.

28. Horcajo P, Ortega-Mora LM, Benavides J, Sánchez-Sánchez R, Amieva R, Collantes-Fernández E, Pastor-Fernández I. 2023. Ovine placental explants: A new ex vivo model to study host-pathogen interactions in reproductive pathogens. Theriogenology 212:157–171.

29. McMillen CM, Arora N, Boyles DA, Albe JR, Kujawa MR, Bonadio JF, Coyne CB, Hartman AL. 2018. Rift Valley fever virus induces fetal demise in Sprague-Dawley rats through direct placental infection. Sci Adv 4:eaau9812.

30. Casazza RL, Lazear HM, Miner JJ. 2020. Protective and Pathogenic Effects of Interferon Signaling During Pregnancy. Viral Immunol 33:3–11.

31. Megli CJ, Coyne CB. 2022. Infections at the maternal-fetal interface: an overview of pathogenesis and defence. Nat Rev Microbiol 20:67–82.

32. Wells AI, Coyne CB. 2018. Type III Interferons in Antiviral Defenses at Barrier Surfaces. Trends Immunol 39:848–858.

33. Ding J, Maxwell A, Adzibolosu N, Hu A, You Y, Liao A, Mor G. 2022. Mechanisms of immune regulation by the placenta: Role of type I interferon and interferon-stimulated genes signaling during pregnancy. Immunol Rev 308:9–24.

34. Koi H, Zhang J, Parry S. 2001. The mechanisms of placental viral infection. Ann N Y Acad Sci 943:148–56.

35. Daubney R, Hudson JR, Garnham PC. 1931. Enzootic hepatitis or rift valley fever. An undescribted virus disease in sheep cattle and man from east africa. Journal of Pathology 34:545–579.

36. Findlay GM, Daubney R. 1931. The virus of Rift Valley fever or enzootic hepatitis. Lancet Glob Health 221:1350–1351.

37. Odendaal L, Clift SJ, Fosgate GT, Davis AS. 2020. Ovine Fetal and Placental Lesions and Cellular Tropism in Natural Rift Valley Fever Virus Infections. Vet Pathol 57:791–806.

38. Robbins JR, Skrzypczynska KM, Zeldovich VB, Kapidzic M, Bakardjiev AI. 2010. Placental syncytiotrophoblast constitutes a major barrier to vertical transmission of Listeria monocytogenes. PLoS Pathog 6:e1000732.

39. Koi H, Zhang J, Makrigiannakis A, Getsios S, MacCalman CD, Strauss JF, 3rd, Parry S. 2002. Syncytiotrophoblast is a barrier to maternal-fetal transmission of herpes simplex virus. Biol Reprod 67:1572–9.

40. Zeldovich VB, Clausen CH, Bradford E, Fletcher DA, Maltepe E, Robbins JR, Bakardjiev AI. 2013. Placental syncytium forms a biophysical barrier against pathogen invasion. PLoS Pathog 9:e1003821.

41. Ander SE, Rudzki EN, Arora N, Sadovsky Y, Coyne CB, Boyle JP. 2018. Human Placental Syncytiotrophoblasts Restrict Toxoplasma gondii Attachment and Replication and Respond to Infection by Producing Immunomodulatory Chemokines. mBio 9.

42. Bayer A, Lennemann NJ, Ouyang Y, Bramley JC, Morosky S, Marques ET, Jr., Cherry S, Sadovsky Y, Coyne CB. 2016. Type III Interferons Produced by Human Placental Trophoblasts Confer Protection against Zika Virus Infection. Cell Host Microbe 19:705–12.

43. Coetzer JA, Barnard BJ. 1977. Hydrops amnii in sheep associated with hydranencephaly and arthrogryposis with wesselsbron disease and rift valley fever viruses as aetiological agents. Onderstepoort J Vet Res 44:119–26.

44. Bird BH, Albariño CG, Nichol ST. 2007. Rift Valley fever virus lacking NSm proteins retains high virulence in vivo and may provide a model of human delayed onset neurologic disease. Virology 362:10–5.

45. Billecocq A, Spiegel M, Vialat P, Kohl A, Weber F, Bouloy M, Haller O. 2004. NSs protein of Rift Valley fever virus blocks interferon production by inhibiting host gene transcription. J Virol 78:9798–806.

46. Koenig ZT. 2018. Type III Interferon Control of Rift Valley Fever Virus at Epithelial Cell Barrier. Master of Science. University of Pittsburgh, SUNY Geneseo.

