## Supplementary material for "Vaccine strains of Rift Valley fever virus exhibit attenuation at the maternal-fetal placental interface": All Supplemental figures

Cynthia McMillen *et al.*

**This PDF file includes:**

Figs. S1 to S3
Tables S1 to S2

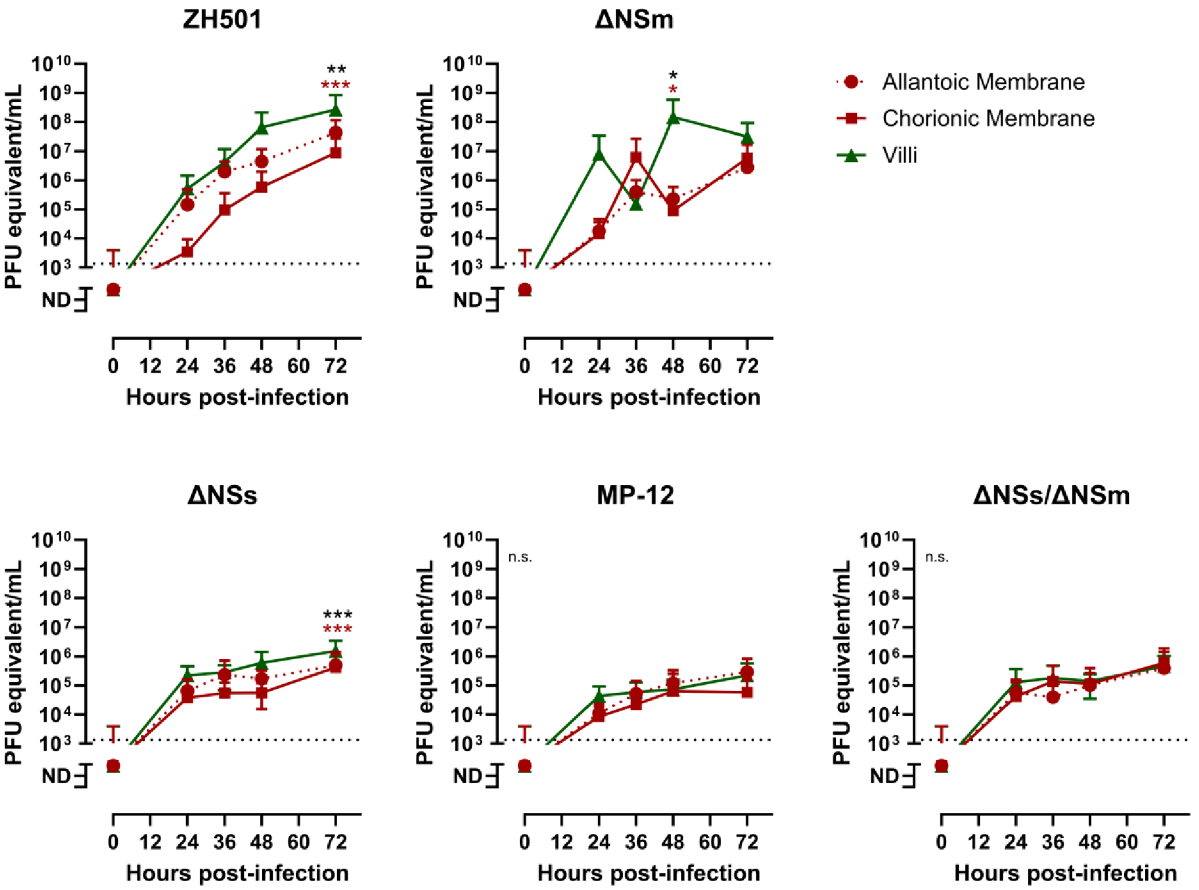


**Fig. S1.** Virus titers within the cultures supernatant at 0, 24, 36, 48, 72 hpi infected with ZH501, ΔNSm (top), ΔNSs, MP-12, ΔNSs/NSm (bottom) compared by sheep placenta tissue type: allantoic membrane, chorionic membrane and villi. Titers were determined by q-RT-PCR. Dashed line = limit of detection (LOD). Statistical significance was determined by a one-way ANOVA compared to ZH501 cultures. * $p<0.05$, ** $p<0.01$, *** $p<0.001$. **** $p<0.0001$. n.s. = not significant. Black asterisks designate the p-values comparing the allantoic membrane and villi. Maroon asterisks designate the p-values comparing the chorionic membrane and the villi.

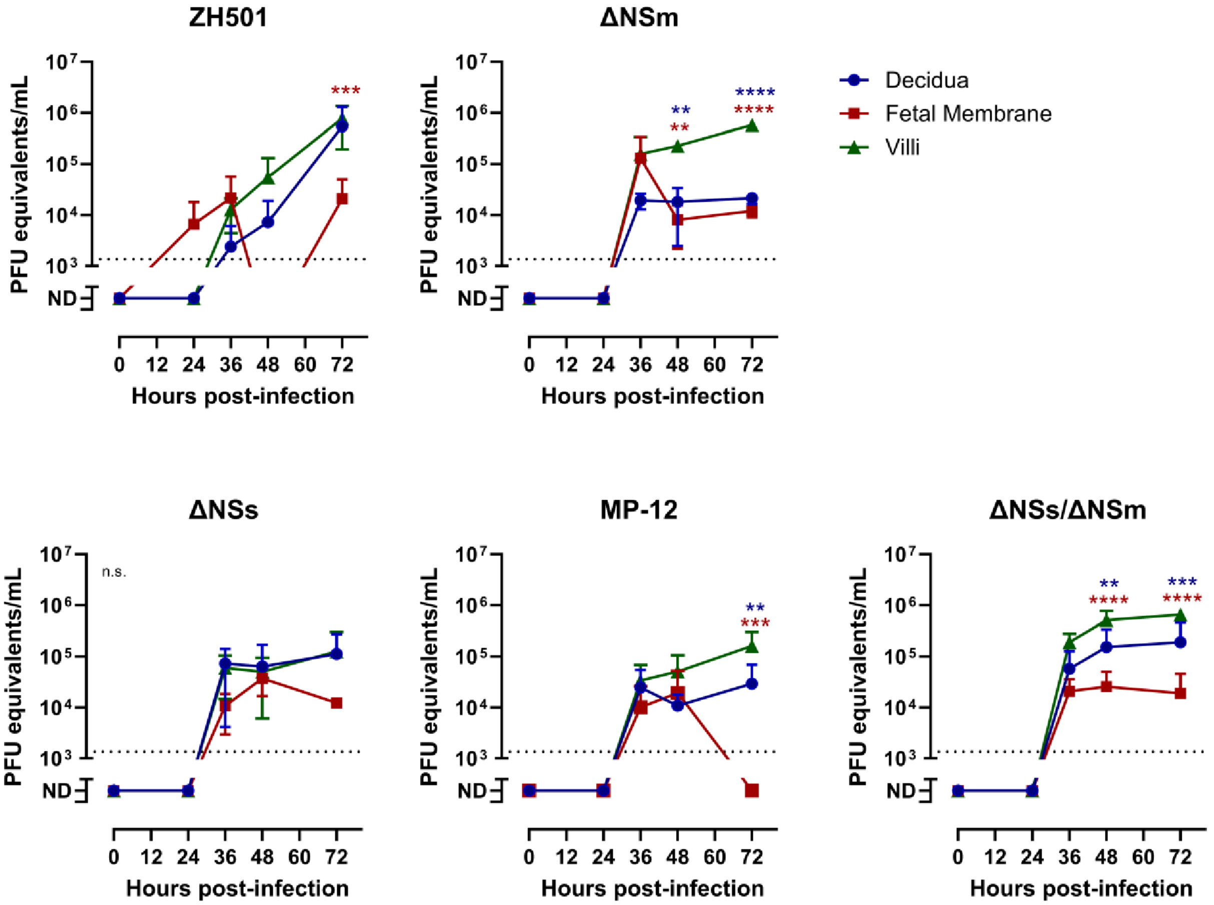


**Fig. S2**. Virus titers within the cultures supernatant at 0, 24, 36, 48, 72 hpi infected with ZH501, ΔNSm (top), ΔNSs, MP-12, ΔNSs/NSm (bottom) compared by human placenta tissue type: decidua, fetal membrane, villi. Titers were determined by q-RT-PCR. Dashed line = limit of detection (LOD). Statistical significance was determined by a one-way ANOVA compared to ZH501 cultures. * $p<0.05$, ** $p<0.01$, *** $p<0.001$. **** $p<0.0001$. n.s. = not significant. Blue or maroon asterisks designate the p-values comparing the decidua or the fetal membrane to the villi, respectively.

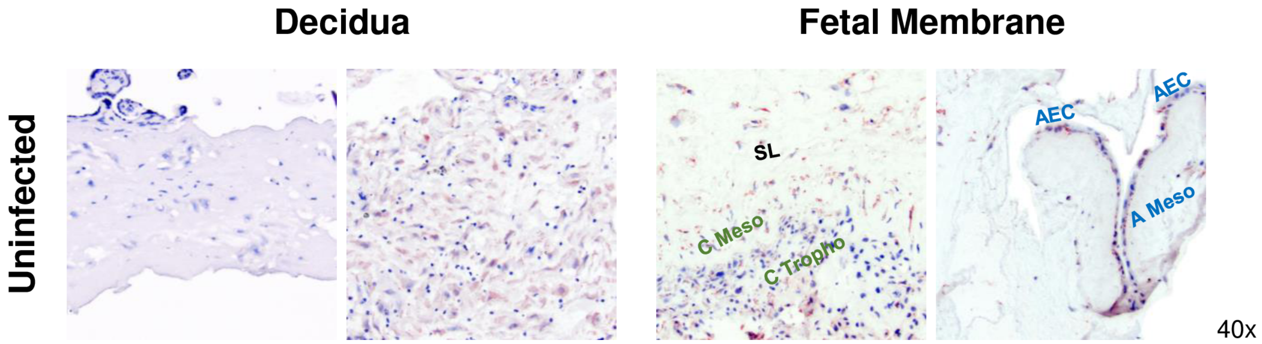


**Fig. S3.** Immunohistochemistry images of uninfected human decidua and fetal membrane cultures. Images were taken at 40x magnification by light microscopy. Blue staining = cell structural counterstain. Structures on the fetal side include the amnion epithelial cells (AEC) and amniotic mesoderm (A Meso). The spongy layer (SL) separates the fetal and maternal sides. Structures on the maternal side include the chorionic mesoderm (C Meso) and the chorionic trophoblasts (C Tropho).

**Table S1. Summary of histological findings in sheep placenta explants.**

| Virus | Tissue | Nucleoprotein Detection (IHC) | | |
|---|---|---|---|---|
| | | *Cellular distribution* | | *Tissue distribution* |
| ZH501 | AM | + (moderate) | cytoplasmic | scattered, in stroma |
| | CM | + (moderate) | cytoplasmic, membranous | |
| | V | + (moderate) | cytoplasmic, membranous | |
| ΔNSm | AM | + (strong) | cytoplasmic, membranous | in epithelium & endothelium |
| | CM | | | |
| | V | + (strong) | cytoplasmic, membranous | |
| ΔNSs – No staining observed | | | | |
| MP-12 | AM | | | |
| | CM | + (moderate) | cytoplasmic, membranous | rarely through stroma & trophoblasts |
| | V | | | |
| ΔNSs/ΔNSm | AM | | | |
| | CM | + (weak) | cytoplasmic | |
| | V | + (strong) | cytoplasmic | scattered in trophoblasts & stroma |
| No infection – No staining observed | | | | |

Staining Intensity Key:

| | |
|---|---|
| + (dark red) | strong |
| + (red) | moderate |
| + (light red) | weak |
| + | intensity not noted |
| | none |

**Table S2. Summary of histological findings in human villous explants.**

| Virus | Nucleoprotein detection (IHC) in villi | |
|---|---|---|
| | Intravillous | Syncytiotrophoblasts |
| ZH501 | + | + |
| ΔNSm | + | + |
| ΔNSs | + | |
| MP-12 | | + |
| ΔNSs/ΔNSm | + | + |
| No infection | | |
